# To Recover or Not to Recover: A Urinary Proteomic Perspective on Severe Acute Kidney Injury

**DOI:** 10.64898/2026.10.12.758641

**Authors:** André Q. Figueiredo, Inês F. Domingos, Sílvia Coelho, Maria G. Cabral, Marta B. Lopes, Luís B. Carvalho, Hugo López-Fernández, José L. Capelo, Carlos Lodeiro, Rajiv Dhir, António Jacinto, Jacek R. Wiśniewski, Gianluca Moroncini, Hugo M. Santos

## Abstract

**Rationale & Objective:** Early prediction of renal non-recovery after acute kidney injury (AKI) could support risk-adapted follow-up and clinical decision-making. We evaluated whether a urinary fatty acid-binding protein 3 (FABP3)/apolipoprotein A-I (APOA1) ratio based on Total Protein Approach (TPA) quantification could discriminate renal recovery from non-recovery after severe AKI.

**Study Design:** Exploratory biomarker discovery study with outcome-blinded independent validation.

**Setting & Participants:** Critically ill adults with KDIGO stage 2 or 3 AKI. Urine was collected 48 hours after AKI diagnosis. The discovery cohort included 20 patients (10 recovered and 10 non-recovered), and the outcome-blinded validation cohort included 10 patients.

**New Predictor:** Urinary TPA-derived FABP3/APOA1 ratio (AKI^TPA index^).

**Outcome:** Renal recovery at day 14, defined as survival with serum creatinine returning to within 0.3 mg/dL or 150% of baseline; patients not meeting these criteria were classified as having non-recovery.

**Analytical Approach:** Urinary proteins were analyzed by LC-MS/MS and quantified using TPA. Differential analysis identified FABP3 and APOA1 as oppositely associated with renal outcome. A predefined FABP3/APOA1 threshold of 1 classified patients as non-recovery-like (>1) or recovery-like (<1) and was subsequently applied to the outcome-blinded validation cohort.

**Results:** In the discovery cohort, the AKI^TPA index^ correctly classified 17 of 20 patients (accuracy, 85.0%; sensitivity, 80.0%; specificity, 90.0%; ROC AUC, 0.930). In the validation cohort, 9 patients were evaluable, of whom 7 were correctly classified (accuracy, 77.8%; sensitivity, 75.0%; specificity, 80.0%; ROC AUC, 0.950).

**Limitations:** The exploratory design and limited cohort size warrant further validation in larger, independent populations.

**Conclusions:** The urinary FABP3/APOA1 TPA ratio showed encouraging preliminary discrimination of renal recovery after severe AKI and warrants prospective clinical validation in larger, independent multicenter cohorts.

---

Acute kidney injury (AKI) is an important risk factor for acute kidney disease (AKD), chronic kidney disease (CKD), and end-stage renal disease and is associated with substantial morbidity, mortality, prolonged hospitalization, and increased healthcare costs ^1,2^. The extent of renal recovery after AKI strongly influences subsequent renal and non-renal outcomes ^3^. However, the conventional markers of AKI, namely serum creatinine (sCr) and urine output, are delayed and relatively insensitive, limiting early diagnosis, risk stratification, and therapeutic decision-making ^4^.

Urinary and plasma biomarkers reflecting tubular injury, metabolic dysfunction, and inflammatory signaling may complement these conventional measures^4,5^. Urinary AKI biomarkers include low-molecular-weight proteins that are normally reabsorbed in the proximal tubule, enzymes released by damaged tubular cells, and proteins upregulated in renal cells after injury ^4,6^. Their combined analysis may improve outcome prediction and provide insight into biological mechanisms underlying AKI ^7–9^. Urinary biomarker dysregulation is often related to the severity and anatomical distribution of tubular damage ^10^.

Proteomics extends disease characterization beyond genomics by measuring dynamic changes in protein abundance, post-translational modifications, and pathway activity ^11–13^. High-resolution mass spectrometry combined with the Total Protein Approach (TPA) has been used to identify biomarkers and potentially druggable targets in renal disease^14,15^.TPA estimates absolute protein abundance by expressing each protein signal relative to the total proteome signal, providing a label-free and standard-free approach^16,17^. Although its reliability depends on proteome coverage and analytical performance, TPA is well suited to exploratory biomarker discovery and longitudinal analysis ^14,15,18,19^.

Among emerging AKI biomarkers, fatty acid-binding protein 3 (FABP3) and apolipoprotein A-I (APOA1) are of particular interest because both are involved in lipid metabolism. FABP-family proteins, apolipoprotein biology, and peroxisome proliferator-activated receptor (PPAR)-related signaling have all been linked to cellular stress, metabolic dysfunction, and tissue injury ^20–24^. FABP3 is a cytoplasmic lipid-binding protein involved in intracellular fatty-acid transport and energy metabolism. Increased FABP3 levels have been associated with renal replacement therapy in AKI and with AKI after cardiac surgery ^20,25^, suggesting that high urinary FABP3 may reflect an injury-dominant phenotype.

APOA1, the major apolipoprotein of high-density lipoprotein (HDL), has been associated with protection against tissue injury and with early renal dysfunction ^21–23^. It is also linked to PPAR-regulated lipid metabolism, which may be relevant to renal tubular biology ^21–23^. However, urinary APOA1 may reflect several processes, including increased glomerular permeability, impaired proximal tubular reabsorption, altered HDL metabolism, or the presence of APOA1-containing lipid particles ^22,26^. Its urinary abundance should therefore not be regarded as a direct surrogate for circulating HDL concentration or function^27^. Within a proteomic classifier, APOA1 may instead reflect a broader biological state involving lipid transport, inflammatory regulation, and preservation or restoration of renal metabolic function.

Ratios between biologically contrasting proteins may be more robust and easier to interpret than individual protein measurements. They can reflect the relative predominance of opposing biological processes while partially reducing the influence of urine concentration, sample loading, and total protein abundance, as previously studied by our group ^28^. Building on this concept, the present study evaluated whether a TPA-based FABP3/APOA1 ratio, referred to as the AKI^TPA index^,, could distinguish renal recovery from non-recovery after AKI. We hypothesized that a ratio greater than 1, indicating FABP3 predominance, would reflect an injury-associated, non-recovery-like phenotype, whereas a ratio below 1, indicating APOA1 predominance, would reflect a more favorable recovery-like phenotype. The ratio was first developed in a discovery cohort and then assessed in an outcome-blinded independent validation cohort. The aim was to determine its discriminative performance, transferability, and clinical interpretability for predicting renal recovery after AKI.

## Methods

### Sample collection and patient inclusion criteria

The cohort used in this study has been described previously ^29^. Briefly, adult patients admitted to the intensive care unit (ICU) with severe AKI, defined as KDIGO stage 2 or 3 based on sCr, were enrolled. At 48 h after AKI diagnosis, 20 mL of freshly voided urine was collected into ethylenediaminetetraacetic acid (EDTA)-containing tubes and centrifuged at 3000 rpm for 5 min to remove cell debris. Finally, the concentrated urinary proteomes were quantified using a Bradford Protein Assay (Sigma-Aldrich), with bovine serum albumin (BSA) used for the calibration curve. The study was conducted in accordance with the protocol approved by the Medical Ethics Review Committee of Hospital Fernando da Fonseca EPE (reference 18.02/2016). Total recovery of renal function was defined as being alive at day 14 with a sCr return to within 0.3 mg/dL or 150% of baseline sCr. Patients who did not meet these criteria were classified as having non-recovered renal function. Written informed consent was obtained from each patient or, when appropriate, from a legally authorized representative. Patients were excluded if they met any of the following criteria: absence of urinary dipstick analysis at ICU admission; anuria; renal replacement therapy during the week preceding ICU admission; established chronic kidney disease exceeding stage 3, as estimated using the CKD-EPI equation in accordance with KDIGO guidelines; kidney, urinary tract, or prostate cancer; ICU length of stay shorter than 48 h; or absence of written informed consent.

The final study population comprised two datasets. The discovery cohort included 20 patients, of whom 10 experienced renal recovery and 10 did not. Each sample was analyzed in two technical replicates, except for one replicate from patient 10 in the non-recovery group, which was excluded from downstream analyses. The outcome-blinded independent validation cohort comprised 10 patients whose clinical outcomes were blinded during proteomic analysis. Each validation sample was analyzed in three technical replicates (Fig. 1). For the validation cohort, outcome data were withheld from the analytical team throughout proteomic processing and interpretation. Clinical outcomes were disclosed by the treating physicians only after patient-level classifications had been generated, thereby ensuring an outcome-blinded validation procedure.

**Figure 1.**
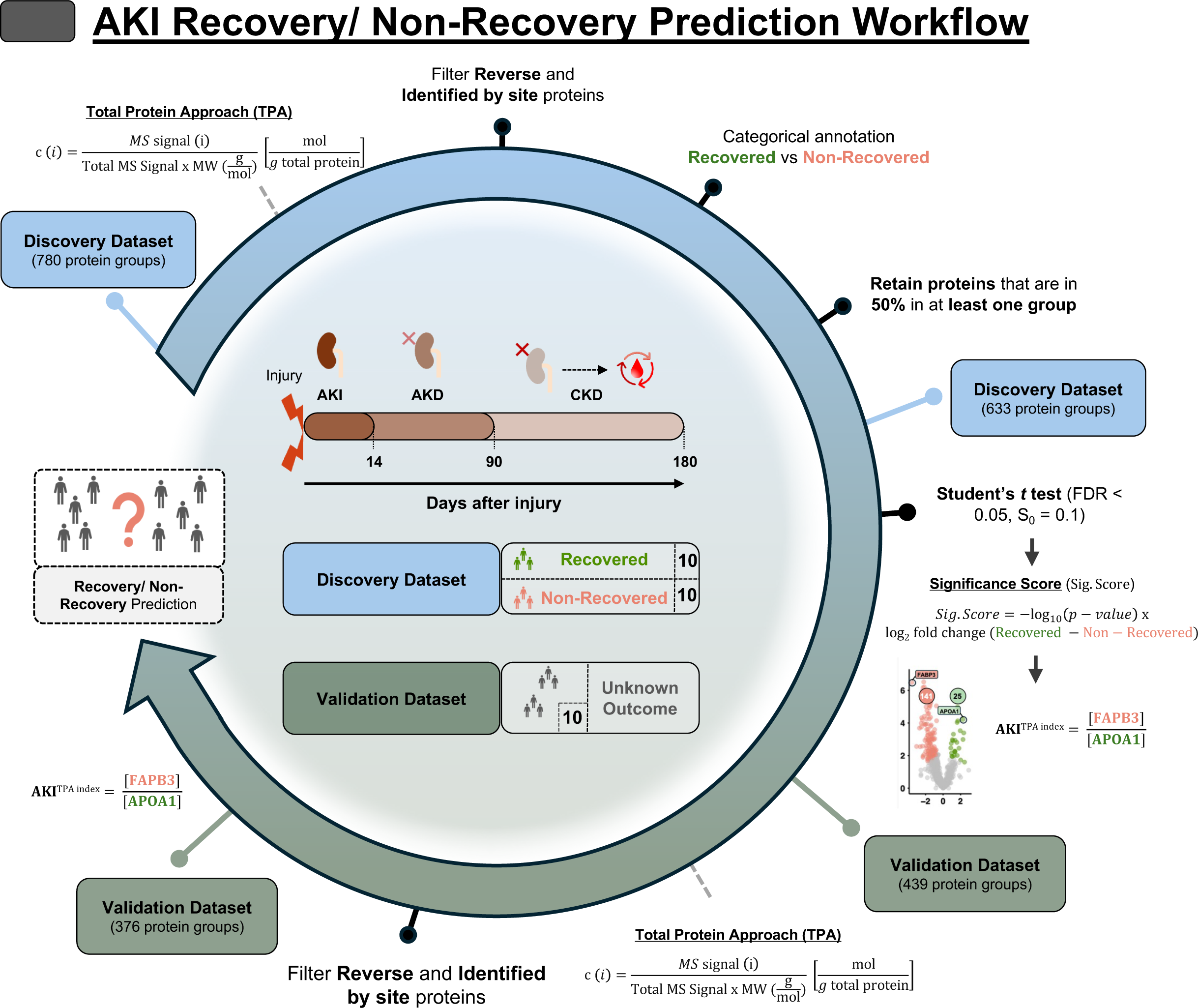
Discovery and validation workflow for a FABP3/APOA1 ratio-based biomarker of acute kidney injury (AKI) recovery prediction. The discovery dataset (*n* = 20) was analyzed using the Total Protein Approach (TPA; Equation 1) to estimate absolute protein concentrations. Reverse hits and proteins identified exclusively by modification site were removed, and samples were classified according to renal outcome as recovery or non-recovery. Proteins detected in at least 50% of technical replicate measurements in either outcome group were retained for differential analysis using a two-sided Student’s *t*-test, with a permutation-based false discovery rate (FDR) of 0.05 and a *S*_0_ parameter of 0.1. Significance scoring (Equation 2) identified FABP3 and APOA1 as oppositely regulated candidate biomarkers. Based on these findings, a threshold-based classifier, termed the AKI^TPA index^, was defined as the ratio of the TPA-derived concentrations of FABP3 and APOA1 (FABP3/APOA1; Equation 3). The outcome-blinded independent validation cohort (*n* = 10) was processed using the same TPA calculation and outcome-independent preprocessing, without repeating discovery-stage outcome-dependent filtering, differential analysis, or biomarker selection used in the discovery dataset. Each validation patient was then classified using the predefined AKI^TPA index^. Ratio values >1 were interpreted as a non-recovery-like profile, whereas values <1 were interpreted as a recovery-like profile.

Urinary proteome preparation and nano-liquid chromatography–tandem mass spectrometry (nLC-MS/MS) analysis are thoroughly explained in Supplementary Material 1.

### Data analysis and bioinformatics workflow

Relative label-free quantification was performed using the precursor signal intensity method and delayed normalization, MaxLFQ, on MaxQuant software V2.0.3.0. All raw files were processed in two separate runs with default parameters^30,31^. Database searches were performed using Andromeda against the human UniProt UP000005640_9606 database. Searches were configured with cysteine carbamidomethylation as a fixed modification and N-terminal acetylation and methionine oxidation as variable modifications. The false discovery rate (FDR) was set to 0.01 for protein and peptide levels with a minimum length of seven amino acids for peptides. The FDR was determined by searching a reverse database. Enzyme specificity was set as C-terminal to arginine and lysine as expected using trypsin. A maximum of two missed cleavages were allowed. Prior to data processing, reverse hits and proteins identified only by site were removed. Then, the Total Protein Approach (TPA) was applied to the protein-group MS Intensity values exported from MaxQuant, rather than MaxLFQ-normalized intensities, to estimate the absolute concentration of each protein *i* in both datasets (Equation 1). The principle of TPA relies on the assumption that the total spectral intensity from all detected peptides in a mass spectrometry (MS)-based experiment is directly proportional to the concentration of the protein in the solution^16^. The protein molecular weight is then used to convert this relative signal into an absolute concentration in molar units (mol of protein per g of total protein):

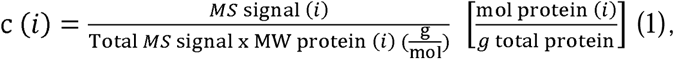

where *MS* signal (i) corresponds to the spectral intensity of protein *i*, the Total *MS* signal is the sum of all protein spectral intensities in each sample, and the MW, the corresponding molecular weight for that protein *i* ^16^.

Consequently, for any given protein with a measurable spectral intensity, a TPA-based concentration was calculated (Supplementary Material 2). Processed TPA-based concentration data were further analyzed using Perseus v2.0.11, as described below^32,33^. TPA-based concentrations were log_2_-transformed to reduce data skewness and the influence of extreme values. Pearson correlation analysis was subsequently performed on log_2_-transformed TPA-based values for both datasets, retaining individual technical replicate measurements. In the discovery cohort, protein groups were retained if detected in at least 50% of replicate measurements in either outcome group, and Z-scored log_2_-transformed TPA-based values were used for unsupervised hierarchical clustering. Differentially abundant proteins between the recovery and non-recovery groups were identified using a two-sided Student’s t-test applied to the individual technical replicate measurements, with a permutation-based false discovery rate (FDR) of 0.05 and *S*_0_ = 0.1 (Supplementary Material 2). For each statistically significant protein, a Significance Score (Sig. Score) that combines both the log_2_ fold change and the -log_10_ (*p*-value) was applied, as *per* Equation 2 (Fig. 1):

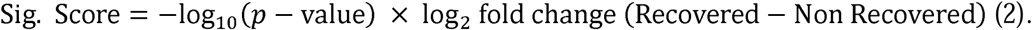

This Sig. Score was calculated using a similar approach as described by Xiao *et al* ^34^. The protein with the highest positive Sig. Score (APOA1, enriched in recovered patients, yielding a positive log_2_ FC when expressed as Recovered – Non-Recovered) and most negative Sig. Score (FABP3, enriched in non-recovered patients, yielding a negative log_2_ FC when expressed as Recovered – Non-Recovered) values were chosen for a TPA-based ratio to predict AKI recovery, labelled as AKI^TPA index^, calculated using the following Equation 3:

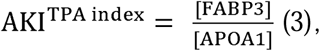

where [FABP3] and [APOA1] are TPA-based concentrations for these proteins in each sample. Technical replicate protein concentrations were averaged at the patient level for ratio calculation and performance assessment. Then, this threshold-based decision rule using the FABP3/APOA1 TPA ratio was applied not only to the discovery dataset but also to an outcome-blinded independent validation cohort to assess its potential for predicting AKI recovery. AKI^TPA index^ > 1 indicated FABP3 dominance and was classified as non-recovery-like, while a value of AKI^TPA index^ < 1 indicated APOA1 dominance and was classified as recovery-like. Values exactly equal to 1 were considered indeterminate because FABP3 and APOA1 were present at equivalent TPA-derived molar concentrations.

### Statistical analysis

Discriminative performance of the AKI^TPA index^ was evaluated separately in the discovery and validation cohorts, with renal non-recovery defined as the positive outcome. Performance measures included accuracy, sensitivity, specificity, positive and negative predictive values. For the described parameters, 95% confidence intervals were calculated using the Wilson score method. Receiver-operating-characteristic (ROC) analysis was used to evaluate the discriminative ability of the continuous AKI^TPA index^, quantified by the area under the ROC curve (ROC AUC).

Patients in whom both FABP3 and APOA1 were undetected were considered analytically non-evaluable because the resulting 0/0 ratio is indeterminate and were therefore excluded from the primary classification metrics. Such patients should be reported separately as analytically indeterminate rather than assigned to either recovery or non-recovery. Because the number of patients and outcome events was limited, the resulting estimates are expected to have substantial uncertainty. No formal claims of clinical validity were therefore made. All statistical analyses were regarded as exploratory and hypothesis-generating. Larger prospective cohorts will be required to obtain more precise estimates of discrimination, assess calibration, determine clinically appropriate decision thresholds and evaluate whether the AKI^TPA index^, provides incremental prognostic value beyond established clinical variables.

## Results

### Proteomics analysis and TPA-based protein profiles

In the discovery dataset, the overall similarity among patient proteomic profiles was evaluated by calculating pairwise Pearson correlation coefficients, which were visualized as a hierarchically clustered correlation matrix (Fig. 2a). Correlation coefficients ranged from 0.44 to 0.99, while replicate measurements showed a mean correlation coefficient of 0.99, indicating excellent technical reproducibility.

**Figure 2.**
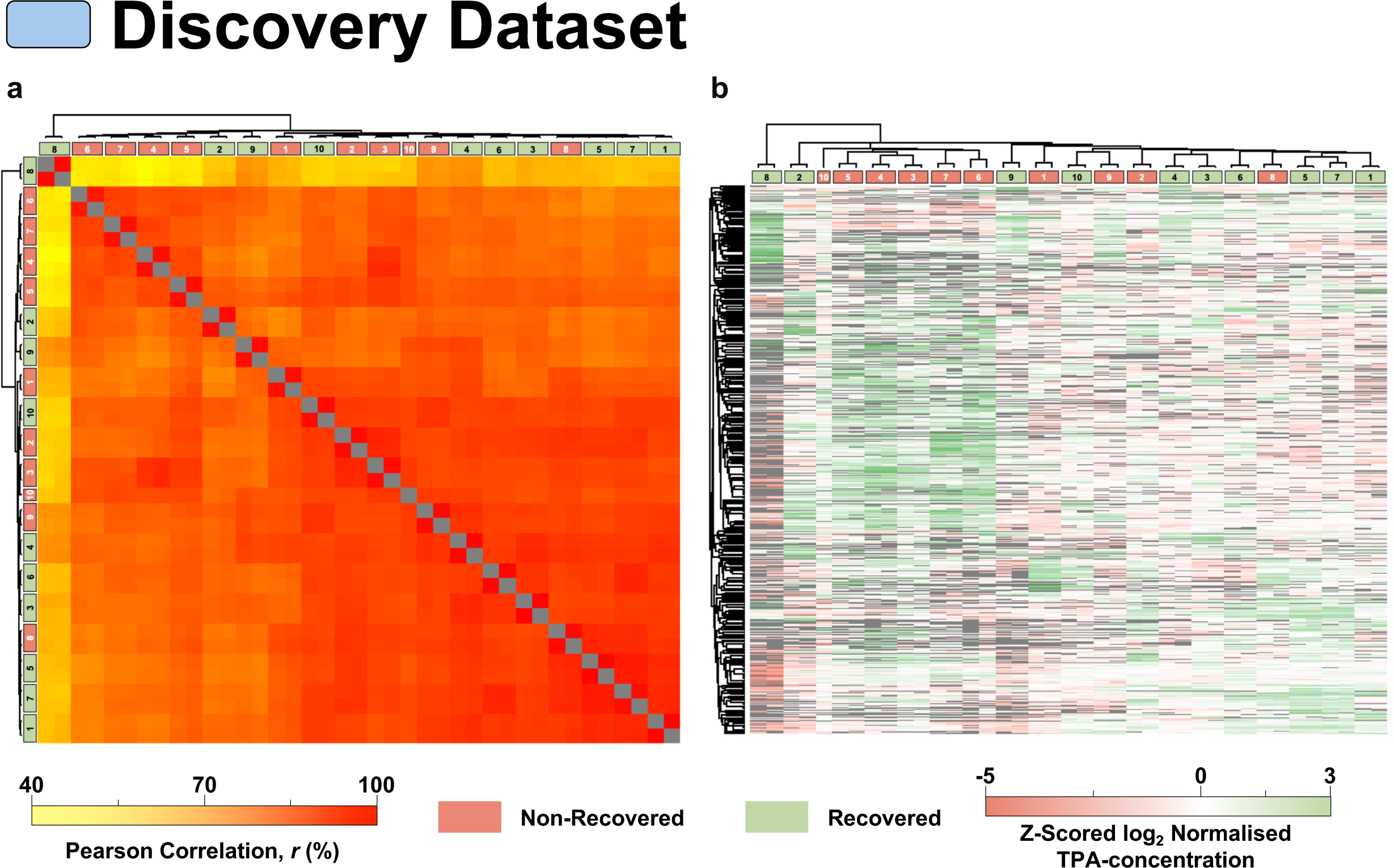
Hierarchical clustering and correlation heatmap of the discovery dataset. **(a)** Pairwise Pearson correlation heatmap of the 39 technical replicate measurements obtained from the 20 discovery patients, with hierarchical clustering applied to both rows and columns. Pearson correlation coefficients are expressed as percentages, and the grey diagonal represents perfect self-correlation (*r* = 1). **(b)** Heatmap of row-wise z-scores calculated from log_2_-transformed TPA-derived protein concentrations across the discovery samples (*n* = 20, 39 technical replicate measurements). Hierarchical clustering was applied to both samples and proteins to identify groups with similar abundance profiles. Light green and light red indicate recovered and non-recovered samples, respectively. Missing values are shown in grey.

To investigate protein abundance patterns in the discovery dataset, hierarchical clustering was performed using Z-scores calculated from log_2_-transformed TPA-derived protein concentrations, and the results were visualized as a heatmap (Fig. 2b). Samples from recovered and non-recovered patients were interspersed across the major sample clusters. Although some protein subsets displayed localized abundance differences and small clusters appeared enriched for a particular clinical outcome, the overall proteomic profiles did not show a clear separation according to recovery status in the discovery cohort.

### Development and patient-level classification of AKI^TPA index^

In the discovery cohort (*n* = 20), differential abundance analysis was performed using TPA-derived protein concentrations to identify proteins associated with renal outcome. The results were visualized in a volcano plot (Fig. 3a). Accordingly, positive fold-change values indicate higher protein abundance in patients who recovered, whereas negative values indicate higher abundance in patients who did not recover.

**Figure 3.**
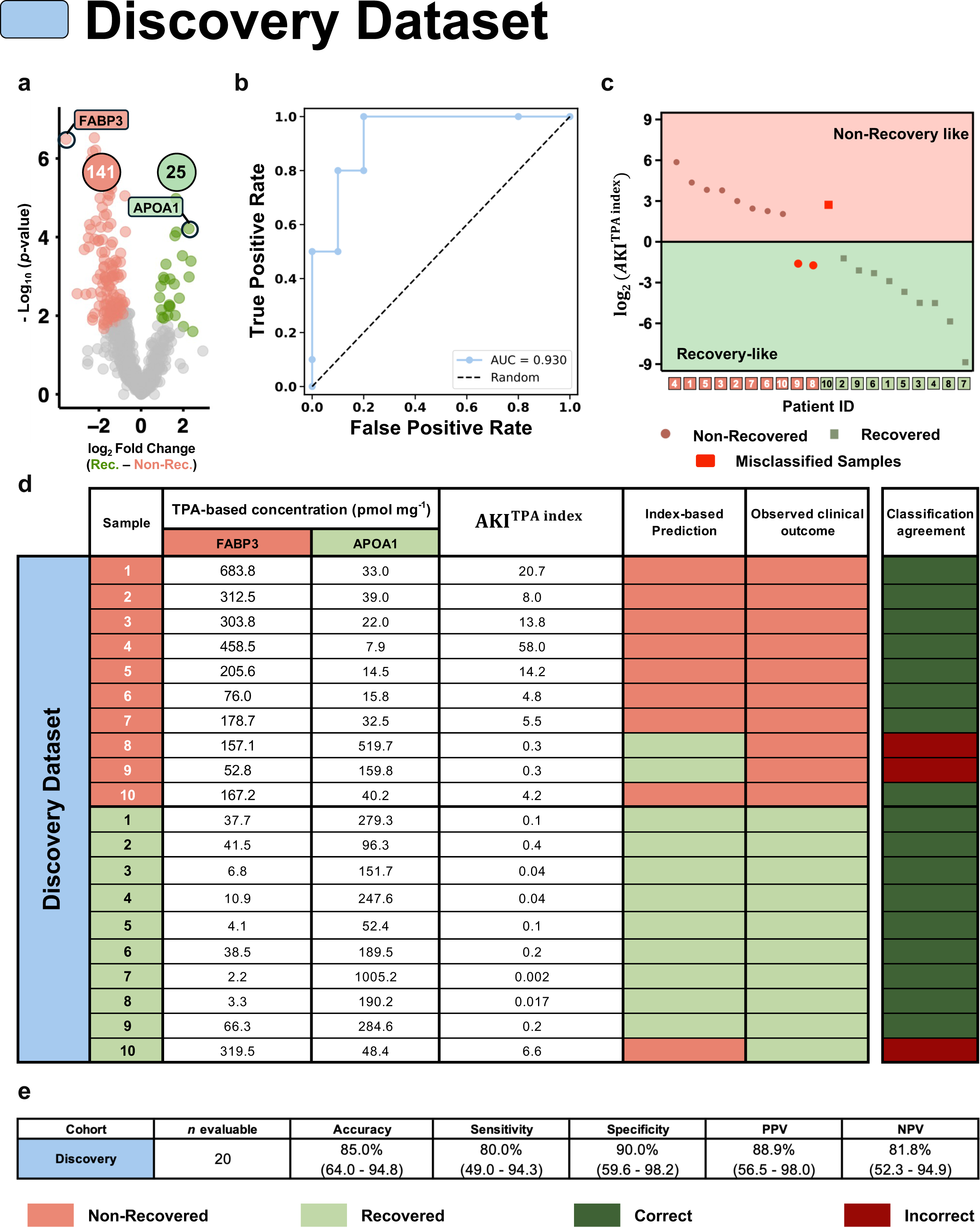
Derivation and apparent performance of the AKI^TPA index^ in the discovery cohort. (**a**) Volcano plot comparing TPA-derived protein concentrations between technical replicate measurements from recovered (*n* = 10) and non-recovered (*n* = 10) patients in the discovery cohort. The x-axis represents the log□ fold change calculated as Recovered/Non-recovered, such that negative values indicate higher abundance in non-recovered patients and positive values indicate higher abundance in recovered patients. Group differences were assessed using a two-sided Student’s *t*-test with a permutation-based false discovery rate (FDR) of 0.05, *S*_0_ = 0.1. A total of 141 proteins showed higher abundance in non-recovered patients, whereas 25 showed higher abundance in recovered patients. FABP3 and APOA1 were selected because they exhibited the most negative and most positive Significance Scores, respectively. (**b**) Receiver operating characteristic curve showing the apparent discriminatory performance of the AKI^TPA index^, defined as the FABP3/APOA1 ratio, in the discovery cohort. Non-recovery was considered the positive outcome. The index yielded an area under the curve of 0.930. Because biomarker selection and performance assessment were conducted in the same cohort, this analysis represents apparent rather than independently validated discrimination. (**c**) Distribution of the logo-transformed AKI^TPA index^ across the 20 patients. Patients are grouped according to their observed clinical outcome and ordered by decreasing index value within each group. Using the cutoff of 1.0, index values > 1.0 [log□(AKI^TPA index^) > 0] were classified as non-recovery-like, whereas values < 1.0 [log□(AKI^TPA index^) <0] were classified as recovery-like. Marker shape indicates the observed clinical outcome (circles, non-recovered; squares, recovered), while the shaded regions indicate the index-based prediction. Misclassified samples are highlighted in red. (**d**) Sample-level results showing the TPA-derived concentrations of FABP3 and APOA1, the calculated AKI^TPA index^, the index-based prediction, the observed clinical outcome, and classification agreement. (**e**) Apparent classification performance of the AKI^TPA index^ at the cutoff of 1.0. Seventeen of the 20 patients were correctly classified, corresponding to an accuracy of 85.0%, sensitivity of 80.0%, specificity of 90.0%, positive predictive value of 88.9%, and negative predictive value of 81.8%. Values in parentheses represent 95% confidence intervals calculated using the Wilson score method. Positive and negative predictive values reflect the balanced outcome distribution of the discovery cohort and should not be interpreted as estimates for populations with a different prevalence of non-recovery.

Using the significance score defined in Equation 2, fatty acid-binding protein 3 (FABP3) exhibited the most pronounced association with non-recovery, whereas apolipoprotein A-I (APOA1) showed the strongest association with recovery. Specifically, FABP3 was more abundant in the non-recovered group, while APOA1 was more abundant in the recovered group. Given these opposing abundance patterns, the ratio of their TPA-derived concentrations was evaluated as a composite outcome-related marker. This ratio, calculated as [FABP3]/[APOA1], was designated the AKI^TPA index^ (Equation 3). The biological plausibility of this inverse relationship is supported by previous reports linking FABP3 with tissue injury and adverse renal outcomes, and APOA1 with lipid transport, metabolic regulation, and protective responses^20,22,25,35^. Ratio values greater than 1 indicate that the urinary TPA-derived concentration normalized to total protein of FABP3 exceeds that of APOA1, with patients classified as a non-recovery-associated profile, whereas values below 1 indicate predominance of APOA1, with patients classified as a recovery-associated profile.

The discriminatory performance of the AKI^TPA index^ was evaluated in the 20 patients included in the discovery cohort using ROC analysis, with an AUC of 0.930 (Fig. 3b), demonstrating strong discrimination between recovered and non-recovered patients. The patient-level distribution of the AKI^TPA index^ on a log_2_ scale is shown in Fig. 3c. Corresponding TPA-derived concentrations of FABP3 and APOA1, together with the resulting AKI^TPA index^, predicted class, clinically observed outcome, and classification agreement (Yes, green; No, red), are presented in Fig. 3d. Three patients were misclassified by the AKI^TPA index^ : patients 8 and 9 from the non-recovered group were incorrectly classified as recovery-like, whereas patient 10 from the recovered group was incorrectly classified as non-recovery-like. The overall discriminative performance of the AKI^TPA index^ is summarized in Fig. 3e. The AKI^TPA index^ correctly classified 17 out of 20 patients according to their true recovery status, yielding an overall accuracy of 85.0% (95% CI, 64.0–94.8%). Sensitivity was 80.0% (95% CI, 49.0–94.3%), with two patients in the non-recovered group incorrectly classified as recovery (false negatives). Specificity was 90.0% (95% CI, 59.6–98.2%), while the positive and negative predictive values were 88.9% (95% CI, 56.5–98.0%) and 81.8% (95% CI, 52.3–94.9%), respectively. Although these findings indicate good overall classification performance in the discovery cohort, the relatively wide confidence intervals reflect the small sample size and indicate substantial uncertainty around the point estimates. Therefore, the observed performance should be interpreted cautiously and validated in larger independent cohorts.

### Classification performance of AKI^TPA index^ in an outcome-blinded independent validation cohort

To evaluate the generalizability of AKI^TPA index^, its discriminative performance was applied to an outcome-blinded independent validation cohort (*n* = 10). Before disclosure of patient outcomes, the AKI^TPA index^ was applied to the independent validation dataset samples to maintain an unbiased analysis, with patient outcomes provided by the physicians only after classification. As in the discovery dataset, Pearson correlation analysis was first performed in the outcome-blinded independent validation cohort to assess replicate and sample concordance (Fig. 4a). Patient 7, with neither FABP3 nor APOA1 detection, was considered analytically indeterminate, preventing calculation of a meaningful ratio. ROC analysis demonstrated good discriminatory performance of the AKI^TPA index^, yielding an AUC of 0.950 (Fig. 4b). The patient-level distribution of the AKI^TPA index^ on a log_2_ scale is shown in Fig. 4c. Corresponding TPA-derived concentrations of FABP3 and APOA1, together with the resulting AKI^TPA index^, predicted class, clinically observed outcome, and classification agreement (Yes, green; No, red), are presented in Fig. 4d. Two patients were misclassified by the AKI^TPA index^: Patient 4 (ratio = 0.9) was incorrectly classified as recovery-like by the binary threshold-based decision rule. On the other hand, patient 5 (ratio = 2.4) was incorrectly classified as non-recovery-like. The overall discriminative performance of the AKI^TPA index^ is summarized in Fig. 4e.

**Figure 4.**
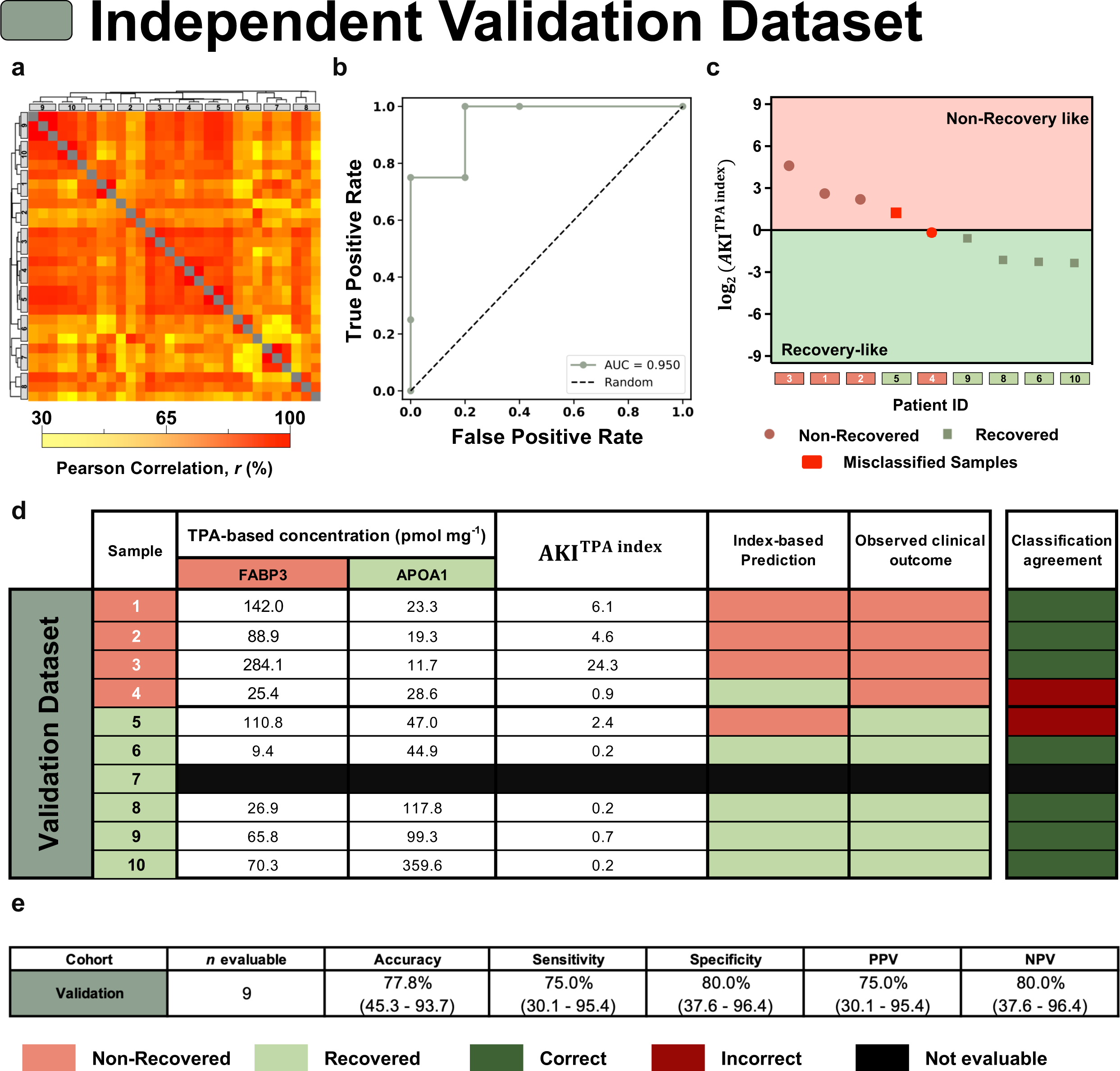
Independent validation of the AKI^TPA index^ in the validation cohort. (**a**) Pairwise Pearson correlation heatmap of the 30 technical replicate measurements obtained from the 10 outcome-blinded validation samples, with hierarchical clustering applied to both rows and columns. Pearson correlation coefficients are expressed as percentages, and the grey diagonal represents perfect self-correlation (*r* = 1). (**b**) Receiver operating characteristic curve for the AKI^TPA index^ (FABP3/APOA1 ratio), in the nine evaluable validation samples comprising four non-recovered and five recovered patients. Non-recovery was defined as the positive outcome. The index yielded an area under the curve of 0.950. (**c**) Distribution of the log□-transformed AKI^TPA index^ among the nine evaluable patients. Using the cutoff of 1.0, index values > 1.0 [log□(AKI^TPA index^) > 0] were classified as non-recovery-like, whereas values < 1.0 [log□(AKI^TPA index^) < 0] were classified as recovery-like. Marker shape indicates the observed clinical outcome (circles, non-recovered; squares, recovered), whereas the shaded regions indicate the index-based prediction. Misclassified samples are highlighted in red. (**d**) Sample-level results showing the TPA-derived concentrations of FABP3 and APOA1, the calculated AKI^TPA index^, the index-based predicted outcome, the observed clinical outcome, and classification agreement. Neither FABP3 nor APOA1 was detected in patient 7, resulting in an undefined 0/0 ratio. Consequently, this patient was considered analytically indeterminate. This patient was retained in the correlation analysis shown in panel a but excluded from the ROC and classification-performance analyses. (**e**) Classification performance of the AKI^TPA index^ at the cutoff of 1.0 in the nine evaluable samples. The index correctly classified seven of nine patients, corresponding to an accuracy of 77.8%, sensitivity of 75.0%, specificity of 80.0%, positive predictive value of 75.0%, and negative predictive value of 80.0%. Values in parentheses represent 95% confidence intervals calculated using the Wilson score method. Positive and negative predictive values are specific to the outcome distribution in this validation cohort.

Among the nine patients evaluated in the outcome-blinded independent validation cohort, the AKI^TPA index^ correctly classified seven according to their true recovery status, yielding an overall accuracy of 77.8% (95% CI, 45.3–93.7%). Sensitivity was 75.0% (95% CI, 30.1–95.4%), with one patient in the non-recovered group (patient 4) incorrectly classified as recovery (false negative). Specificity was 80.0% (95% CI, 37.6–96.4%), while the positive and negative predictive values were 75.0% (95% CI, 30.1–95.4%) and 80.0% (95% CI, 37.6–96.4%), respectively. Although these findings support the classification performance of the AKI^TPA index^ in an independent cohort, the wide confidence intervals reflect the small number of evaluable patients and indicate substantial uncertainty around the point estimates. Therefore, these performance estimates should be interpreted cautiously and require confirmation in larger independent cohorts.

The absence of both proteins in one validation patient illustrates a second clinically relevant outcome: an assay can be technically valid at the global proteome level yet remain non-informative for a specific two-marker threshold-based decision rule using FABP3 and APOA1. In future clinical implementations, such samples could require repeat analysis, targeted measurement, or interpretation using complementary clinical variables.

## Discussion

The present analysis identifies the urinary AKI^TPA index^, derived from the TPA-based FABP3/APOA1 ratio, as a promising and clinically interpretable decision rule for renal recovery after severe acute kidney injury. The index correctly classified 17 of 20 patients in the discovery cohort and 7 of 9 evaluable patients in the outcome-blinded independent validation cohort. Its fixed threshold of 1 provides a straightforward interpretation: values >1 indicate that FABP3 is more abundant than APOA1 and are classified as a FABP3-dominant, non-recovery-like pattern, whereas values <1 indicate APOA1 predominance and a recovery-like pattern. Importantly, although FABP3 and APOA1 were selected based on their differential abundance in the discovery cohort, the threshold itself was not data-derived or optimized to maximize classification accuracy. Rather, it was predefined at a ratio of 1, corresponding to equal TPA-derived molar concentrations of FABP3 and APOA1. This biologically interpretable cutoff may facilitate translation by providing a simple molecular decision rule without requiring a complex predictive model.

The biological direction of the ratio is also plausible. FABP3 is involved in intracellular long-chain fatty-acid transport and cellular energy metabolism. Increased extracellular concentrations can reflect membrane disruption, metabolic stress or tissue injury. Previous studies have associated urinary FABP3 with severe AKI and the need for renal replacement therapy, and perioperative FABP3 has also been linked to AKI after cardiac surgery^20,25^. Its association with non-recovery may therefore reflect persistent injury and metabolic dysfunction. Proximal tubular cells depend strongly on mitochondrial oxidative phosphorylation and fatty-acid oxidation, both of which are disrupted during severe AKI^8^. Persistent mitochondrial dysfunction, impaired fatty-acid oxidation and lipid accumulation can contribute to energetic failure, oxidative stress and maladaptive repair^36,37^, potentially producing a FABP3-dominant urinary pattern. Nevertheless, urinary FABP3 may not be kidney-specific, because cardiac or skeletal-muscle injury could also increase circulating FABP3 and influence urinary abundance^35,38^.

APOA1, in contrast, is the major structural and functional apolipoprotein of HDL and contributes to reverse cholesterol transport, antioxidant activity, inflammatory regulation and endothelial protection^39,40^. These functions are relevant to AKI, in which oxidative stress, endothelial activation, leukocyte recruitment and microvascular injury influence persistence and recovery^36,37^. Experimental studies support a potentially protective role for APOA1-related pathways: in renal ischemia–reperfusion, APOA1 administration reduced renal dysfunction and inflammatory responses^41^. Urinary APOA1 has also been reported as an early biomarker of AKI^21^. A low AKI^TPA index^ may indicate that APOA1-associated lipid-transport or anti-inflammatory biology is relatively preserved, or simply that FABP3-associated injury is less dominant. This interpretation remains a mechanistic hypothesis rather than evidence of causality. Furthermore, the ratio should not be considered a direct measure of circulating HDL concentration or HDL function^27^.

Prediction of renal recovery remains an important unmet need. Clinical assessment still relies mainly on serum creatinine, urine output, dialysis requirement and overall clinical trajectory^4^, variables that provide limited information about the underlying balance between injury and repair. Several urinary recovery-associated biomarkers have been investigated, but findings remain inconsistent and no single marker has achieved sufficient reliability for routine clinical use^42^. Recent proteomic studies have further shown that AKI comprises molecularly distinct sub phenotypes and that urinary protein profiles differ according to recovery trajectory^43,44^. The present study extends this concept by reducing a broader proteomic signal to a simple two-protein balance.

The AKI^TPA index^ should therefore be considered an exploratory prognostic tool that complements, rather than replaces, clinical assessment. Larger multicenter prospective studies are required to validate the threshold across different AKI etiologies and severities, assess longitudinal changes, and evaluate outcomes including short-term recovery, dialysis independence, 90-day kidney function and CKD progression. Future studies should also determine whether the ratio improves discrimination, calibration, reclassification and decision-curve net benefit beyond established variables such as age, baseline kidney function, AKI stage, etiology, urine output and dialysis requirement.

In conclusion, the AKI^TPA index^ provides a simple, biologically interpretable urinary proteomic approach for distinguishing recovery-from non-recovery-associated patterns after severe AKI.

Although these findings remain exploratory and require confirmation in larger prospective cohorts, the AKI^TPA index^ represents a promising approach for supporting prognostic assessment of renal recovery after severe AKI.

## Data Statement

The mass spectrometry proteomics data have been deposited to the ProteomeXchange Consortium via the PRIDE^45^, at http://www.ebi.ac.uk/pride, partner repository with the dataset identifier PXD081917.

## Acknowledgements

This work was supported by national funds from FCT – Fundação para a Ciência e a Tecnologia (FCT/MCTES), I.P., through projects UID/50006/2025, UID/PRR/50006/2025, UIDB/50006/2020, UIDP/50006/2020, and LA/P/0008/2020 of the Associated Laboratory for Green Chemistry – LAQV REQUIMTE, UID/297/2025 and UID/PRR/297/2025 of the Center for Mathematics and Applications – NOVA Math, and UID/00667/2025 of UNIDEMI. The work was also supported by the FCT project “Peptide Boolean Biosensors: Integrating Proteomics, Computing, and Biosensing for Next-Generation Diagnostics” (2024.13101.PEX). H.M. Santos acknowledges his research contract funded under LA/P/0008/2020. A.Q.F. is supported by FCT PhD grant 2023.00528.BD, and I.F.D. by FCT/MCTES PhD grant 2024.00745.BD. PROTEOMASS Scientific Society is acknowledged for support provided to the Laboratory for Biological Mass Spectrometry – Isabel Moura through funding programmes #PM001/2025, #PM001/2024, #PM001/2019, and #PM003/2016. This publication is based upon work from COST Action PerMediK CA21165, supported by COST (European Cooperation in Science and Technology).

## Authors’ contributions

AQF, LBC, JLC, and HMS designed the study, performed the statistical analysis and developed the concept. JLC, CL, HMS, RD, AJ, and GM provided financial support. SC, MGC, and AJ collected the samples and provided the clinical support. LBC, IFD, and AQF performed the laboratory work under the supervision of JLC and HMS. AQF, IFD, LBC, and JLC drafted the manuscript. MBL, SC, MGC, CL, RD, JRW, AJ, GM, and HMS revised the drafted version, corrected it, and provided valuable suggestions. All authors have approved the final version of the manuscript.

