## Supplementary Material 1 for "To Recover or Not to Recover: A Urinary Proteomic Perspective on Severe Acute Kidney Injury"

### **Supplementary Methods**

#### **Filter-Aided Sample Preparation (FASP)**

Urinary proteome preparation for both datasets was performed using filter-aided sample preparation (FASP), as previously described^15^, with some modifications. The supernatant obtained after the initial centrifugation was loaded onto a 10 kDa molecular-weight cut-off (MWCO) centrifugal filter and centrifuged at 14,000 × *g* for 20 min at 25 °C. Sample loading was continued until 25 µg of total urinary protein had been retained on the membrane. Proteins were reduced with tris(2-carboxyethyl)phosphine (TCEP) and alkylated with chloroacetamide (CAA), followed by centrifugation under the same conditions. The retained proteins were then washed sequentially with Tris-HCl and triethylammonium bicarbonate (TEAB), using the same centrifugal settings. Sequencing-grade trypsin prepared in TEAB buffer at pH 8.0 was added to the membrane, and proteolytic digestion was performed overnight at 37 °C. The following day, the filter units were transferred to clean collection tubes, and peptides were recovered with TEAB, followed by two additional elution steps with 3% (v/v) acetonitrile (ACN) containing 0.1% (v/v) formic acid (FA). Each elution was followed by centrifugation at 14,000 × *g* for 10 min at 25 °C. Finally, the recovered peptide solutions were transferred to 500 µL microcentrifuge tubes, dried to completeness, and stored at −20 °C until nano-liquid chromatography–tandem mass spectrometry (nLC-MS/MS) analysis.

## **LC‒MS/MS**

LC-MS/MS analysis was performed using an UltiMate 3000 ultra-high-performance liquid chromatograph (Thermo Scientific) coupled to an Ultra High-Resolution Quadrupole Time-of-Flight (UHR-QTOF) IMPACT HD mass spectrometer (Bruker). A 3 μL aliquot of sample, at a total peptide concentration of 0.1 μg/μL, was loaded onto a μPAC™ trapping column and desalted for 2.7 min with 1% (v/v) acetonitrile in 0.1% formic acid at a flow rate of 15 μL/min. Peptides were then separated on an analytical 200 cm μPAC™ PharmaFluidics column using a linear gradient at 500 nL/min, with mobile phase A consisting of 0.1% (v/v) formic acid and mobile phase B consisting of 99.9% (v/v) acetonitrile and 0.1% (v/v) formic acid. The gradient was programmed as follows: 0–2 min, 3–5% B; 5–90 min, 5–35% B; and 90–95 min, 35–86% B. Chromatographic separation was performed at 35 °C. MS acquisition was carried out in MS mode (2 Hz) followed by MS/MS acquisition (8–32 Hz), with a total cycle time of 3.0 s, active exclusion, exclusion after one spectrum, and release after 2 min. Precursors were reconsidered for fragmentation if their current intensity was at least 3-fold higher than the previous intensity, with a fragmentation threshold of 2500 counts. All generated mass-spectrometry proteomics data were submitted to the ProteomeXchange Consortium^33^ and are accessible through the PRIDE^34^ repository under the identifier PXD081917.
